# Computational design of Small Transcription Activating RNAs (STARs) for versatile and dynamic gene regulation

**DOI:** 10.1101/169391

**Authors:** James Chappell, Alexandra Westbrook, Matthew Verosloff, Julius B. Lucks

## Abstract

A longstanding goal of synthetic biology has been the programmable control of cellular functions. Central to this goal is the creation of versatile regulatory toolsets that allow for programmable control of gene expression. Of the many regulatory molecules available, RNA regulators offer the intriguing possibility of *de novo* design – allowing for the bottom-up molecular-level design of genetic control systems. Here we present a computational design approach for the creation of a bacterial regulator called Small Transcription Activating RNAs (STARs) and create a library of high-performing and orthogonal STARs that achieve up to ∼9000-fold gene activation. We then demonstrate the versatility of RNA-based transcription control by showing the broad utility of STARs – from acting synergistically with existing constitutive and inducible regulators, to reprogramming cellular phenotypes and controlling multigene metabolic pathway expression. Finally, we combine these new STARs with themselves and CRISPRi transcriptional repressors to deliver new types of RNA-based genetic circuitry that allow for sophisticated and temporal control of gene expression.

## Introduction

RNA is increasingly recognized as a powerful biomolecule for controlling gene expression and engineering synthetic cellular functions^1-4^. One of the reasons for this is that naturally abundant RNA-based regulators and numerous engineered versions are now available to control almost every aspect of gene expression^5^. In addition these regulatory functions can be enacted and tuned by the programmable formation of specific RNA structures, which mediate interactions with cellular machineries to perform gene regulation. For example in bacteria, the formation of simple RNA structures such as hairpins within mRNAs can prevent gene expression machinery from both transcribing and translating mRNAs^2^. Moreover, these *cis*-acting RNA structures can be further controlled through interacting with *trans*-acting small RNAs (sRNAs) or binding of a ligand to prevent or allow their formation, in effect creating inducible genetic control elements^2,3^. This combination of versatile genetic regulation controlled by simple RNA structures creates the intriguing possibility of using nucleic acid design algorithms to create a comprehensive toolset of RNA regulatory elements *de novo*^1,2^. Thus RNA as a substrate for versatile molecular programming has a potential major advantage over less designable protein-based genetic control, and there is great promise for RNA synthetic biology to allow for the bottom up molecular-level design of genetic control systems.

Recently, there has been great progress towards this vision, with numerous demonstrations of synthetic RNA regulators that have been rationally engineered from natural versions^6-11^ or designed *de novo*^12-21^. However, while computational design of RNA regulators has been possible for many years, a major challenge has been the design of high-performing mechanisms that exert large fold changes in gene expression comparable to protein-based regulators. This is recently beginning to change, most notably for RNA-based translational regulators. For example, improvements of biophysical models of translation initiation have resulted in more accurate design of constitutive^22^ and ligand switchable^23^ *cis*-acting translational regulators. In addition, there has been tremendous advances in the *de novo* design of orthogonal libraries of *trans*-acting sRNA translational activators called toehold switches that function with dynamic ranges in the hundreds of fold activation^21^. This has been possible through the combination of powerful RNA design algorithms with design motifs for translational regulation based upon the conditional formation of simple RNA hairpins that are particularly amenable to computational design.

While significant advances have been made in the computational design of translational RNA regulators, it has been more difficult to design RNA regulators of transcription. This is an important challenge, as RNAs that control transcription have the potential to be more general, able to regulate both RNA synthesis and downstream protein synthesis. In addition, they also offer the distinct advantage of being able to implement genetic regulatory circuits entirely as RNAs^7^ that operate on the fast timescales of RNA degradation^24^. The challenge of the *de novo* design of RNA transcriptional regulators is due to several reasons. First, these regulators depend on the formation of transient RNA structures such as intrinsic terminator hairpins that form cotranscriptionally in a kinetically-driven, out-of-equilibrium folding regime^25-28^. However, RNA folding and design algorithms have largely been developed to model and design RNA structures that are in an equilibrium folding regime. Thus RNA design algorithms are not naturally suited to the design of transient RNA structures. Second, the RNA structures underpinning natural switchable transcriptional RNA regulators such as riboswitches are often more complex than just simple hairpins, and depend on interactions that are harder to design such as pseudoknots and non-canonical base pairs^29^. While there has been some progress in designing riboswitches^15,16^, these challenges have so far prevented the *de novo* design of sRNA transcriptional regulators that show protein-like dynamic ranges.

One solution to this challenge is to identify transcriptional regulatory mechanisms that function through simple RNA structures that use motifs that design algorithms can exploit. Towards this end, we recently created synthetic sRNA transcriptional activators called Small Transcription Activating RNAs (STARs) (**Figure 1a**), which are based upon the conditional formation of simple RNA hairpins^30^. This regulator is composed of a target RNA placed upstream of a gene to be regulated that, once transcribed, folds into an intrinsic terminator hairpin that prevents transcription of the downstream gene (OFF state). To activate transcription, a STAR is expressed that interacts with the target RNA, preventing terminator formation and allowing transcription of the downstream gene (ON state). While our original development of STARs provided a simple transcriptional activation mechanism, our approach was dependent on the unpredictable reengineering of natural terminator hairpin sequences into STARs. This resulted in a small number of orthogonal STARs and often low fold of activation, which could only be modestly improved through RNA engineering strategies^31^.

**Figure 1.**
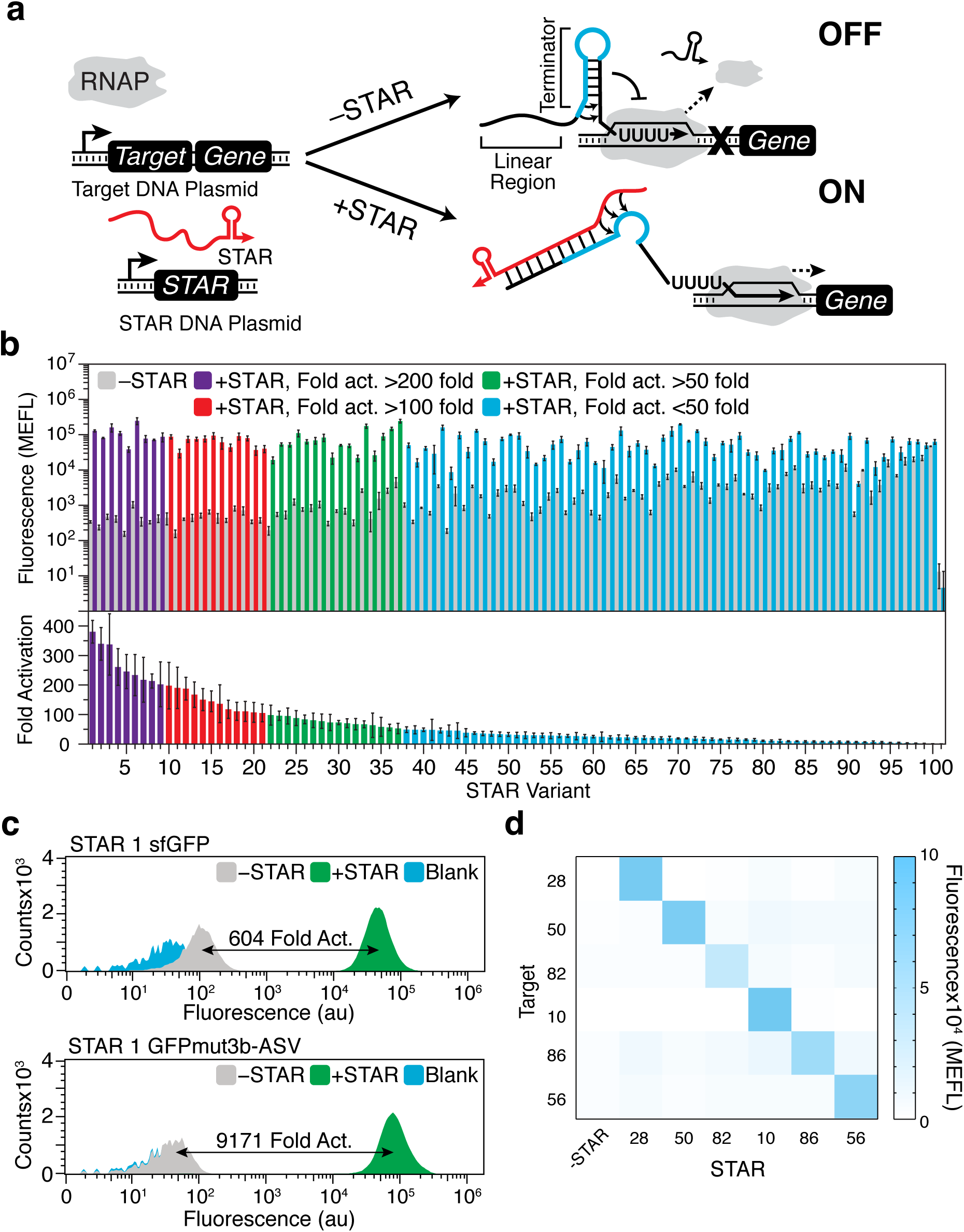
Characterization of a computationally designed small transcription activating RNA (STAR) library. (**a**) Schematic of the small transcription activating RNA (STAR) mechanism. A STAR target DNA sequence is placed upstream of a gene to be regulated. The target RNA is designed to fold into an intrinsic transcription terminator hairpin, composed of a hairpin structure followed immediately by a poly uracil sequence. The formation of this terminator hairpin causes RNA polymerase (RNAP) to terminate transcription upstream of the gene to be regulated (gene OFF). STARs (colored red) bind to both the linear region and the 5’ half of the terminator hairpin (colored blue) of the target RNA, preventing terminator formation and allowing transcription elongation of the downstream gene (gene ON). (**b**) Characterization of 100 computationally designed STAR/target variants and the original AD1 STAR (variant 28)^30^. Fluorescence characterization was performed on *E. coli* transformed with a plasmid encoding each target RNA variant controlling super folder GFP (sfGFP) expression in the absence (-STAR) and presence (+STAR) of a plasmid encoding its cognate STAR. Upper panel shows mean fluorescence and lower panel shows fold activation (ON/OFF), both colored accordingly to the fold activation. A Welch’s *t*-test was performed on each -STAR/+STAR condition with all STAR variants showing statistically significant differences between -STAR and +STAR conditions (*P* < 0.05). We note that variant 101 showed no activation. (**c**) Flow cytometry histograms of two STAR variants expressing sfGFP and GFPmut3b-ASV in the absence (-STAR) and presence (+STAR) of cognate STAR, compared to the autofluorescence of *E. coli* cells transformed with control plasmids (Blank). (**d**) Characterization of 6 STAR variants predicted to be orthogonal (**Supplementary Fig. 10**) by challenging target RNAs controlling sfGFP expression against cognate and non-cognate STARs. Fluorescence characterization was performed as in (**b**) with *E. coli* cells transformed with each STAR/target combination. Cognate pairs are across the diagonal. Mean fluorescence in the absence of STAR (-STAR) is shown in the left column with raw data shown in **Supplementary Fig. 11**. Data were measured with flow cytometry in units of arbitrary fluorescence (au, panel **c**) or Molecules of Equivalent Fluorescein (MEFL, panel **b, d**). Data in (**b**,**d**) represent mean values of at least *n* = 7 biological replicas ± s.d. and (**c**) a representative flow cytometry histogram of *n* = 1 biological replicas with repeats shown in **Supplementary Fig. 6**.

However, the inherent simplicity of the STAR mechanism could potentially permit the application of nucleic acid design algorithms for the *de novo* design of STARs to address these current limitations. To deliver on this potential, we describe here the creation of a new STAR design motif that allowed for the computational design of large libraries of STARs with high-dynamic ranges. Moreover, our computational design approach offered an immediate solution to the often complex problem of creating independently acting, or orthogonal, RNA regulators. This was achieved through computational screening to identify STAR/target pairs with minimal non-cognate STAR-target interactions followed by experimental validation of these computational predictions.

These achievements then allowed us to demonstrate the versatility and power of designable RNA transcriptional control in a number of contexts of importance to synthetic biology approaches and applications. Because STARs control transcription, they can be easily interfaced with other commonly used constitutive and inducible regulatory systems to tune their performance. Specifically, we show that STARs are capable of interfacing with libraries of constitutive promoters and ribosome binding sites to create switchable versions of these elements, which have themselves been mainstays of synthetic biology. We also show that the performance of inducible protein-based promoter systems can be tuned by interfacing them with a STAR without the need for protein or protein-DNA interaction engineering. We next demonstrate that these STARs can function robustly in different genetic and environmental contexts by showing that they can control diverse messenger RNAs *in vivo* encoded on plasmids and the genome, and can tightly control gene expression within *in vitro* cell-free gene expression systems. We also show broad utility of these STARs – from reprogramming cellular phenotypes to controlling metabolic pathway expression. We anticipate this will have an immediate impact for a range of existing applications from strain and metabolic pathway engineering, to cell-free based diagnostics^32,33^.

Finally, we demonstrate several contexts in which STARs can be leveraged to create RNA-only genetic network motifs of increasing sophistication. This includes the creation of one-to-many control of gene expression with a STAR single-input module, the creation of an activation-activation STAR cascade, and the creation of a split STAR AND gate. Furthermore, interfacing the complementary regulation offered by both STARs and CRISPR interference (CRISPRi) transcriptional repressors^34^ allowed construction of an activation-repression cascade and a NIMPLY logic-gate, which when combined enable us to create the first RNA-based incoherent feed forward loop that creates a pulse of gene expression.

## Results

### Computational design of orthogonal STAR libraries with high-dynamic ranges

As a starting point towards computationally designing STARs, we first sought to uncover a simple STAR design motif by mechanistically analyzing the role of each sequence region of the STAR-target complex (**Supplementary Fig. 1**). Through this analysis we observed that while STAR sequestration of the terminator hairpin is required to prevent terminator formation and activate transcription, STAR binding of the linear region appeared to be critical for seeding this interaction. As such, we inferred that the interaction between the linear region of the target RNA and corresponding binding region of the STAR serves as the key recognition region of the STAR regulatory system. This observation raised the intriguing possibility that functionally diverse and orthogonal STARs could be simply created by varying the linear region while keeping constant the terminator scaffold of the most efficient AD1 STAR system^30^.

We next sought to use this design motif to create a library of STAR variants using the Nucleic Acids Package (NUPACK) RNA online design tool^35,36^ (**Supplementary Fig. 2** and **Supplementary Note 1**). NUPACK is a suite of algorithms that perform statistical thermodynamic calculations of RNA structures to either predict an RNA structure from a given sequence, or to design RNA sequences that will fold into user-defined structures. While NUPACK is a powerful suite of design tools, its application to designing STARs is nontrivial. This is because the statistical thermodynamic calculations behind NUPACK’s design and prediction algorithms are focused on RNA structures that are at equilibrium, whereas STAR regulation is likely governed by kinetic, out-of-equilibrium folding regimes. Nevertheless, by focusing the design on the STAR-target linear recognition region while maintaining the terminator scaffold we hypothesized that we could utilize NUPACK to design efficient interaction sequences that limited competing intramolecular structures, while maximizing intermolecular interactions (**Supplementary Fig. 2**).

Using our STAR design motif and NUPACK, we designed and created a library of 100 STAR variants. To functionally characterize this library, plasmids encoding target RNA-reporter fusions were transformed into *E. coli* cells and fluorescence measured in the presence of plasmids encoding the cognate STAR or a no-STAR control plasmid (**Supplementary Fig. 3**). Characterization of this STAR library revealed a broad-range of functionality, with each STAR showing distinct OFF and ON levels, and folds of activation (**Figure 1b** and **Supplementary Fig. 4**), which we subsequently confirmed were due to changes in transcription using RT-qPCR (**Supplementary Fig. 5**). We observed that 37 variants show > 50 fold activation – comparable to that of the previously best performing STAR^30^ (labeled variant 28 in **Figure 1b**). In addition, numerous variants showed significantly higher folds of activation up to ∼400 fold (**Figure 1c**). Furthermore, we were able to achieve 9171±3881 fold activation by combining the best STAR variant with a commonly used unstable GFP (GFPmut3b-ASV) that reduced the OFF-level fluorescence to be nearly indistinguishable from cellular autofluorescence (**Figure 1c**). Finally, we showed that our best performing STAR compared favorably to the best performing computationally designed toehold switch translational activator^21^, with the distinct advantage of offering lower OFF levels in our experimental configuration (**Supplementary Fig. 6**). As such we showed that we can not only computationally design STARS, but in doing so attain some of the highest-dynamic ranges achieved by an RNA-based regulator to date.

We next aimed to determine whether we could uncover design principles from this functionally diverse STAR library. Studying the functional characteristics of the library first, we observed that a prerequisite for a STAR-target to achieve a high-dynamic range was tight control of gene expression in the OFF state. Thus the transcription termination efficiency of the target RNA represented a key determinant of overall performance. Computational analysis of the library revealed several sequence and structural features that suggested the formation of base stacking interactions and secondary structures within the linear region of the target RNA negatively impacted transcription termination efficiency (see **Supplementary Note 2** for detailed discussion). Thus the best performing STARs more closely satisfied the design criterion of having unstructured linear regions.

We next assessed whether these observations represented a generalizable STAR design principle or were specific for the AD1 terminator hairpin that was used as a terminator scaffold. To test this, we used NUPACK to computationally design a small library of STARs using the terminator from the *E. coli ribA* gene as a terminator scaffold^30^. This ribA STAR library was characterized and shown to be functional, demonstrating that our STAR design strategy was generalizable to other terminator hairpins (**Supplementary Fig. 7**). Interestingly we again observed a strong negative correlation between termination efficiency and the presence of base stacking interactions and predicted secondary structure in the linear region, suggesting this to be general design principle of STARs (see **Supplementary Note 2**).

Finally, we sought to explore whether this design strategy can yield orthogonal, or independently-acting, STAR variants. To do so, we developed a computational algorithm to predict the orthogonality between members of the STAR library (**Supplementary Note 3**). In this algorithm, NUPACK^37^ was first used to predict base pairing between all target RNA and STAR variants (10,201 combinations) (**Supplementary Fig. 8**). This data was then used to select predicted orthogonal subsets of STAR-target variants, by identifying different variants to be orthogonal if the number of predicted base pairs was less than a cut-off value of 13 that we experimentally determined (**Supplementary Fig. 9**). To validate these predictions, we selected a subset of 6 STAR variants that were predicted to be orthogonal (**Supplementary Fig. 10**) and experimentally confirmed that these STARs were indeed highly-orthogonal (**Figure 1d** and **Supplementary Fig. 11**). This demonstrated that orthogonal STARs can be easily identified with purely computational methods, circumventing the need to experimentally screen thousands of possible combinations.

In summary, by combining nucleic acid design algorithms with a simple design motif we can computationally design and create *de novo* large libraries of STARs. Moreover, we showed that through the creation of this library we can identify high-performing and orthogonal variants, and also obtain a deeper understanding of the sequence-structure-function relationship of this regulatory system.

### Interfacing STARs with constitutive and inducible control elements to switch and tune regulatory properties

Synthetic biology has created a wealth of tools for fine-tuning and optimizing multi-gene genetic programs^38^. Perhaps the most prevalent are strength variants of constitutive promoters and ribosome binding sites (RBS). We hypothesized that STARs could be synergistically combined with these constitutive control elements to create switchable version, in which the STAR system provides tight-control of the OFF state and promoter/RBS variants provide tunable control of the ON state. To test this concept, we combined a library of promoter and RBS strength variants (**Supplementary Fig. 12**) with a target RNA expression cassette (**Figure 2a**). Characterization revealed that through different combinations or promoter/RBS variants, highly-tunable activation levels can be achieved – all while maintaining stringent repression in the OFF state (**Figure 2a** and **Supplementary Fig. 13**). Thus STARs provide a generalizable approach to create switchable control elements from the large number of existing libraries of constitutive regulatory parts^39^.

**Figure 2.**
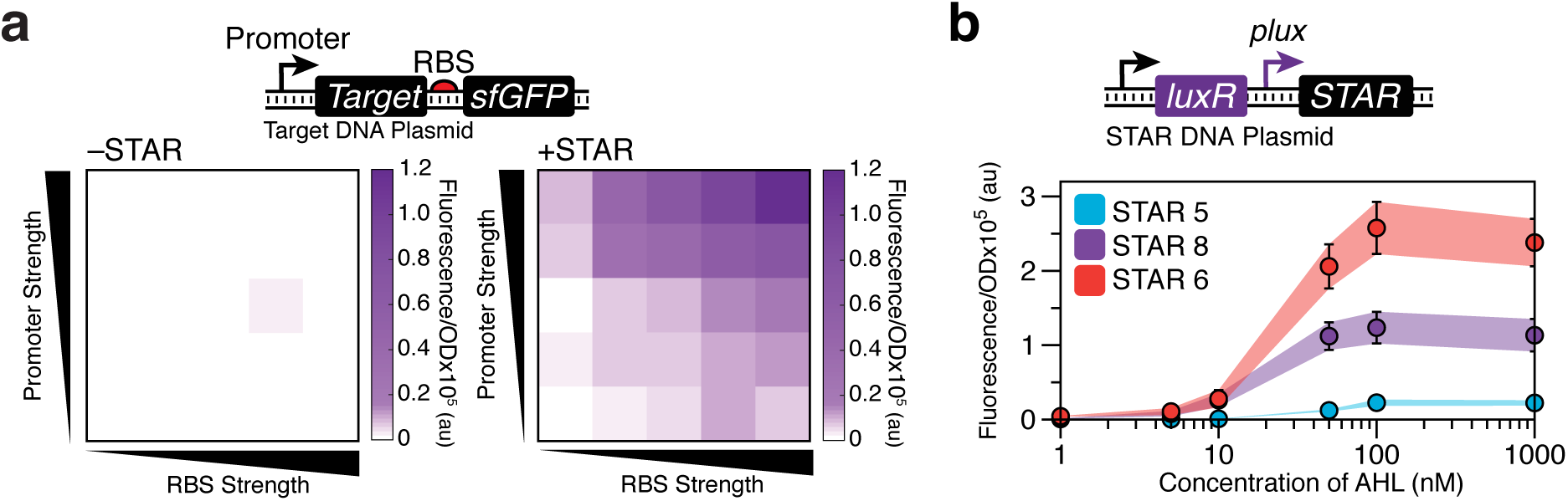
STARs can interface with constitutive and inducible control elements to switch and tune regulatory properties. (**a**) STARs make constitutive gene expression elements switchable. Schematic DNA template of a library of target DNA plasmids containing variable strength promoters and ribosome binding sites (RBS). Fluorescence characterization of *E. coli* cells transformed with this library in the absence (left grid, -STAR) and presence (right grid, +STAR) of a plasmid encoding cognate STAR (raw data shown in **Supplementary Fig. 13**). (**b**) STARs tune inducible promoters. Schematic DNA template of a plasmid in which the *plux* promoter controls STAR production in response to acyl-homoserine lactone (AHL). Fluorescence characterization of *E. coli* cells transformed with this plasmid and a plasmid encoding a target RNA construct controlling sfGFP expression. Changing the STAR/target pair in this configuration results in unique induction curves. Fluorescence characterization was performed by bulk fluorescence measurements (measured in units of fluorescence [FL]/optical density [OD] at 600 nm). Data represents mean values of *n* = 9 biological replicas ± s.d.

The ability of STARs to interface with constitutive control elements motivated us to examine the effects of interfacing STARs with inducible regulators. We first demonstrated that tunable activation could be achieved by using a chemically inducible promoter system to titrate STAR expression and consequently, the level of activation (**Figure 2b**). Interestingly, we also observed that different STAR systems with distinct ON levels resulted in inducible systems with distinct transfer functions, varying in the maximum ON level of expression. Moreover, this ON level could be further fine-tuned through combining this inducible-STAR system with the target-RBS library described above (**Supplementary Fig. 14**). Thus we were able to simply tune the dynamic range of the inducible system without having to undertake the laborious task of either engineering the promoter-transcription factor interface, or the transcription factor itself.

These results demonstrated the power of interfacing easily designable RNA regulators with existing regulatory systems to switch and tune their regulatory properties, all without having to engineer the DNA or protein components.

### STARs function robustly in diverse genetic and environmental contexts

A prerequisite for a robust regulator is that it should predictably function in a variety of contexts. To demonstrate this for STARs, we first confirmed that STAR functionality is maintained independent of the gene being regulated. Three specific STARs were chosen that achieved high-dynamic ranges through a high ON level (STAR 6), low OFF level (STAR 5) or a combination (STAR 8). We combined these STARs to control the expression of three sequence diverse fluorescent proteins and confirmed that the different STARs functioned consistently, independent of the gene being regulated (**Figure 3a**).

**Figure 3.**
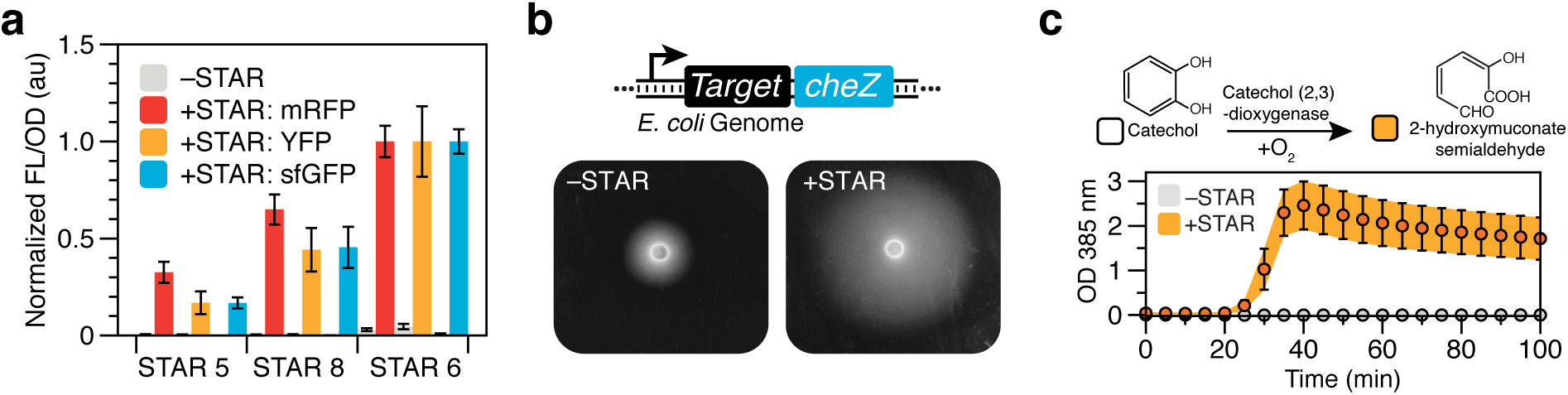
STARs control gene expression tightly in diverse contexts. (**a**) STARs function in the context of sequence diverse fluorescent reporter genes. Fluorescence characterization was performed on *E. coli* cells transformed separately with plasmids encoding three target RNAs controlling sfGFP, YFP or mRFP in the absence (-STAR) and presence (+STAR) of plasmids encoding cognate STARs. Measurements were normalized to 1 for the fluorescence value of STAR variant 6 in the +STAR condition for each fluorescent protein. Fluorescence characterization was performed by bulk fluorescence measurements (measured in units of fluorescence [FL]/optical density [OD] at 600 nm). (**b**) STARs function in the genome to control endogenous *E. coli* genes. Schematic DNA region of *E. coli* BW25113Dc*heZ* modified to contain a genomic copy of a STAR regulated *cheZ* gene. Photographs of semi-solid agar motility assays in the absence (left panel, -STAR) and presence (right panel, +STAR) of a plasmid encoding the cognate STAR. Motility is indicated by the formation of halos around the inoculation site in the center of the photographs. (**c**) STARs function in cell-free transcription-translation (TX-TL) reactions to tightly control enzyme expression. Schematic of the catechol (2,3)-dioxygenase catalyzed reaction that converts colorless catechol into a yellow-colored 2-hydroxymuconate semialdehyde. Spectral characterization of observed yellow color (measured in units of OD at 385 nm) of STAR controlled catechol (2,3)-dioxygenase expression in TX-TL reactions supplemented with catechol in the absence (-STAR) and presence (+STAR) of a plasmid encoding cognate STAR. Data in (**a**, **c**) represents mean values of *n* = 9 biological replicas ± s.d and (**b**) is a representative photograph of *n* = 1 biological replicas with repeats shown in **Supplementary Fig. 16**.

We next demonstrated that STARs can regulate genes located on the genome by integrating a STAR-controlled sfGFP expression cassette into the genome of *E. coli*. This led to a near infinite dynamic range because of the indistinguishable OFF level from cellular autofluorescence (**Supplementary Fig. 15)**. We also demonstrated that STARs could regulate endogenous genes on the genome by engineering a strain of *E. coli* to only express the chemotactic regulator CheZ in the presence of a STAR. As CheZ controls chemotaxis in *E. coli*, we inferred CheZ expression by performing semi-solid agar motility assays. We observed a significant increase in motility in the presence of the cognate STAR, providing further evidence that STARs can control expression of genomic genes, and can even be used to reprogram cellular phenotype (**Figure 3b** and **Supplementary Fig. 16**).

Finally, we demonstrated the ability of STARs to control gene expression within *E. coli* based cell-free transcription and translation (TX-TL) systems^40,41^, which are gaining momentum for a range of applications including as a platform for rapid prototyping of genetic circuitry^24,42^ and paper based diagnostics^32,33^. We demonstrated STARs can tightly regulate the synthesis of both sfGFP (**Supplementary Fig. 17**) and the enzyme catechol 2,3-dioxygenase (C23DO) (**Figure 3c**), all while maintaining their high-dynamic ranges observed *in vivo*. The C23DO regulation example could be particularly useful for alternative colorimetric outputs for diagnostic applications^32,33^.

Taken together, we have shown that STARs function robustly to provide large dynamic range control of gene expression in a wide-variety of genetic and environmental contexts.

### STARs allow control of multi-gene metabolic pathways

Metabolic engineering is a major application of synthetic biology and there has been much interest in applying genetic control strategies to maximize metabolic pathway productivity and yields. Towards this, we investigated how STARs can be used to control the expression of a multi-gene pathway for the production of deoxyviolacein – a purple compound with valuable anti-bacterial, anti-viral, and anti-tumor properties (**Figure 4a**). To do this, a target RNA with a low OFF level was constructed upstream of the four gene deoxyviolacein pathway, and deoxyviolacein production characterized through solvent extraction and spectral quantification (**Figure 4b**). This characterization revealed that STARs allowed for tight control of the deoxyviolacein pathway expression, with almost no deoxyviolacein production observed in the absence of STAR. This result confirmed STARs can tightly control metabolic pathways, and more generally can be used to regulate multi-gene operon expression through a single regulatory locus.

**Figure 4.**
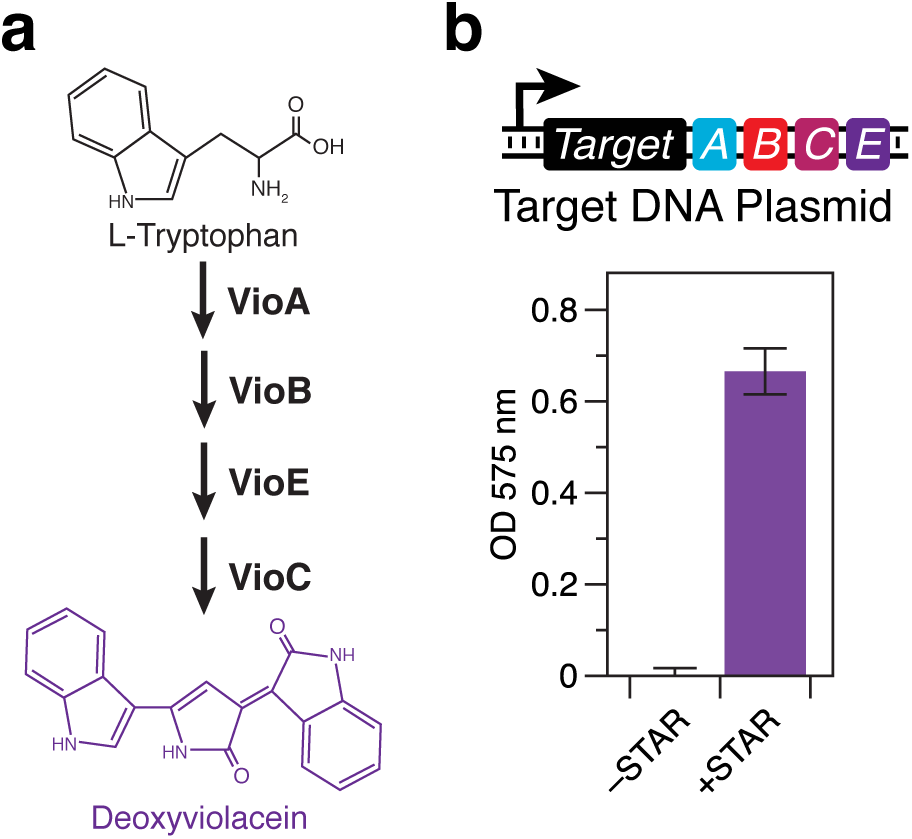
STARs allow for control of multi-gene metabolic pathways. (**a**) Schematic of the four gene deoxyviolacein metabolic pathway (*vioABCE*) that converts L-tryptophan into the purple compound deoxyviolacein. (**b**) Schematic DNA template of a plasmid containing a STAR regulated *vioABCE* pathway. Characterization was performed on *E. coli* DH5 alpha pir cells transformed with this plasmid in the absence (-STAR) and presence (+STAR) of a plasmid encoding cognate STAR. Deoxyviolacein production was characterized by solvent extraction and spectral quantification of deoxyviolacein (measured in units of OD at 575 nm). Data represents mean values of *n* = 9 biological replicas ± s.d.

### STARs allow for the creation of new RNA regulatory network motifs

Construction of synthetic regulatory networks analogous to those seen in natural systems has been a long-standing goal of synthetic biology^43,44^. Towards this goal, we aimed to utilize STARs for creating RNA-only regulatory network motifs in which signals are propagated through transcriptional regulation. We first aimed to use STARs to create a single-input module (SIM)^45^, in which a STAR is used to coordinate expression of multiple genes. In this manner, instead of a STAR controlling a single gene (one-toone) in a cell, a STAR is used to control the expression of multiple genes simultaneously (one-to-many). To test this concept, we configured a high-performing target RNA with a low OFF state upstream of three genes encoding mRFP, sfGFP and the enzyme C23DO. STAR activation of each gene was characterized in cells containing only a single gene at a time (one-to-one), as well as in cells containing all three genes at once (one-to-many) (see **Supplementary Fig.18** for details of C23DO characterization). Characterization revealed the SIM to be functional and able to significantly activate all three genes simultaneously (**Figure 5a**). Interestingly, when the one-to-many and one-to-one circuits were compared, a decrease in the sfGFP expression level in the presence of the STAR was observed in the one-to-many case. We hypothesized that this decrease could be due to either a retroactivity constraint of using the same pool of STAR to activate multiple targets^46^, or from contextual effects of expressing both mRFP and sfGFP from a single plasmid^47^.

**Figure 5.**
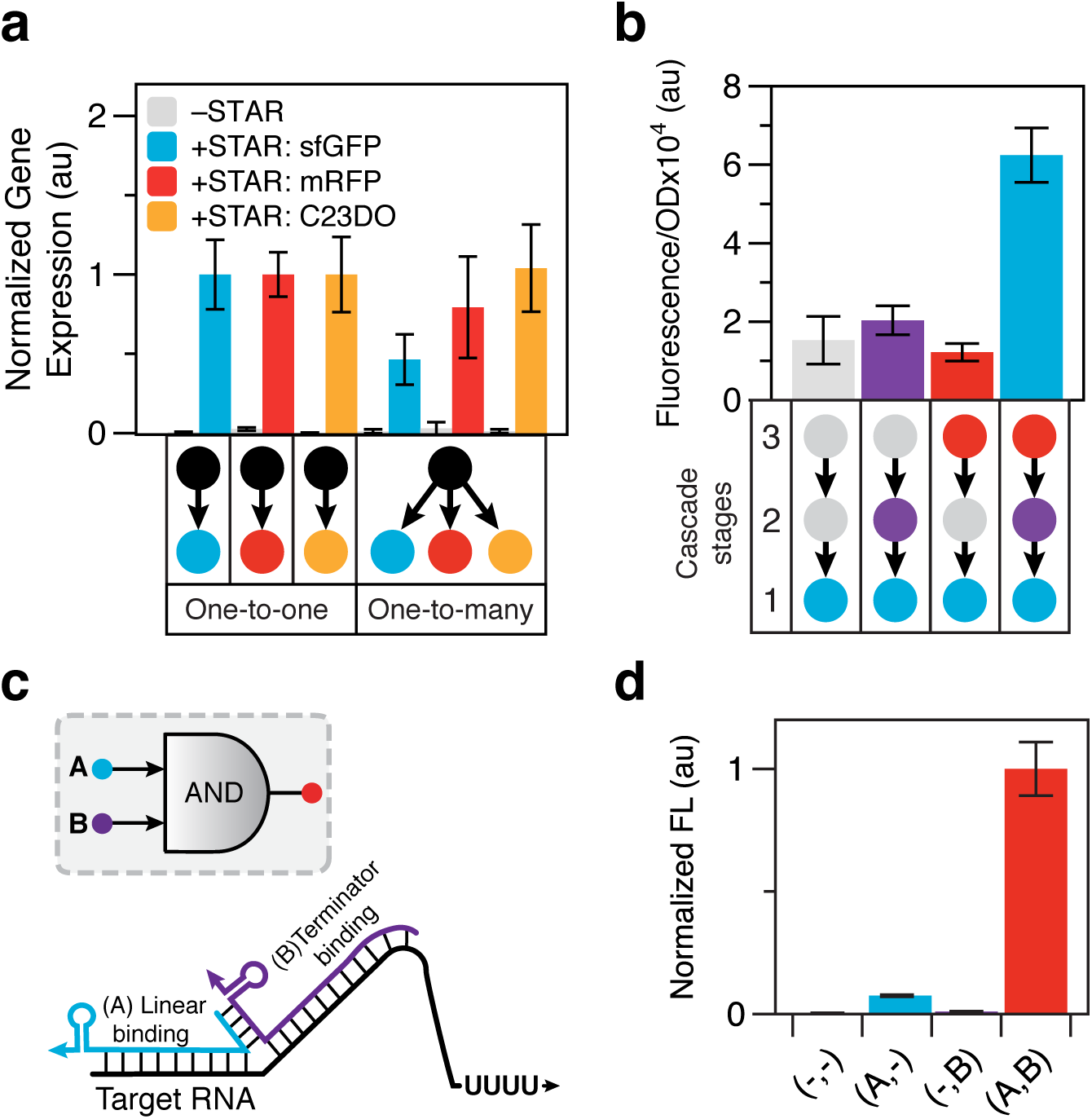
STARs allow for RNA-only network motifs. (**a**) A STAR single-input module (SIM) for one-to-many gene regulation. Fluorescence and spectral characterization of *E. coli* cells transformed with DNA plasmids encoding target RNAs controlling sfGFP, mRFP or catechol (2,3)-dioxygenase (C23DO) expression in the absence (-STAR) and presence (+STAR) of a plasmid encoding the cognate STAR. One-to-one characterization was performed on three different cell strains transformed with one of the plasmids encoding a single STAR regulated gene with and without the STAR plasmid. One-to-many characterization was performed on a single strain transformed with two plasmids encoding all three STAR regulated genes simultaneously with and without the STAR plasmid. C23DO expression level was determined by calculating the maximum rate of conversion of catechol to 2-hydroxymuconate semialdehyde as described in **Supplementary Fig. 18**. Measurements for each gene were normalized to 1 for the one-to-one value for that gene. (**b**) An RNA-only STAR activation-activation cascade. Fluorescence characterization was performed on *E. coli* cells transformed with combinations of DNA plasmids encoding cascade stages or control constructs. Cascade stage combinations used are shown in the schematic below the graph with grey indicating the use of a control DNA plasmid in that condition. Cascade stages were configured so that the bottom (blue) node represents a target RNA controlling sfGFP expression, the middle (purple) node represents a target RNA controlling the synthesis of an orthogonal STAR, and the top (red) node represents a constitutively expressed STAR (see **Supplementary Fig. 19** for plasmid architecture). (**c**) Schematic of a split STAR AND gate. STAR 1 was split into two regions corresponding to the linear and the terminator hairpin binding regions of the STAR (referred to as A and B), with a designed interaction sequence to promote the assembly of the full STAR complex (AB) when both are present. (**d**) Fluorescence characterization was performed on *E. coli* cells transformed with a DNA plasmid encoding a target RNA construct controlling sfGFP expression in the presence of a DNA plasmid encoding different combinations of A and B. Measurements were normalized to 1 for the AB condition. (**a**-**b**) Fluorescence characterization was performed by bulk fluorescence measurements (measured in units of fluorescence [FL]/optical density [OD] at 600 nm) and (**a**) spectral characterization. Raw data for the (**a**) spectral characterization of C23DO production is shown in **Supplementary Fig.18**. (**d**) Fluorescence characterization was performed by flow cytometry (measured in units of Molecules of Equivalent Fluorescein [MEFL]). Data represents mean values of *n* = 9 biological replicas ± s.d.

We next sought to demonstrate that STARs can be used to construct an activation-activation cascade, whereby the transcriptional output of one STAR regulator is used to drive the expression of another STAR. Using two STARs from our orthogonal library, we constructed a three-stage STAR activation-activation cascade. In this design, a STAR activates a target RNA that is transcriptionally fused to an orthogonal STAR, which in turn activates the expression of an sfGFP output (**Figure 5b** and **Supplementary Fig.19**). An endoribonuclease RNA processing platform^48^ was used to separate the transcriptionally fused target RNA-STAR transcript of stage 2 of the cascade. Each of the cascade stages was constructed on separate plasmids and combinations of cascade stages were experimentally characterized (**Figure 5b**). This revealed the cascade to be functional, showing maximal activation (∼4 fold) only in the presence of the full cascade. We note that the fold of activation of the full cascade was lower than that of the lowest fold activation STAR/target pair which limits cascade performance (4 fold compared to 34 fold), which we hypothesized could be due to changes in the relative plasmid copy number encoding the STAR/target RNA.

Finally, we aimed to use STARs to construct an AND logic gate (A AND B). Previously we had shown that this could be achieved by transcriptionally fusing two orthogonal target RNAs in tandem^30^ to perform signal integration at the level of the target RNA. While successful, our mechanistic studies suggested a potentially simpler strategy was to use a single target RNA and perform signal integration at the level of the STAR. This is similar to a recently published strategy with split toehold translational activators^49^. We hypothesized that if a STAR was split into two halves between the linear and terminator binding regions (referred to as A and B), that neither half alone could activate transcription. However, if an interaction sequence was designed between these two STAR halves, when both were present they would form a complex (AB) that would be capable of activating transcription (**Figure 5c**). To test this concept, NUPACK was used to design the interaction sequence between the split STARs. Characterization revealed this AND logic gate design to be functional, only maximally expressing when both STAR halves A and B were present.

In summary we have demonstrated that STARs can be configured to expand the variety of RNA-only transcriptional network motifs that serve as the basic building blocks for the construction of entirely synthetic regulatory networks^45^.

### Combining STARs with CRISPRi to create novel network motifs for temporal control of gene expression

CRISPRi has emerged as a powerful tool to repress transcription in bacteria^34^, while STARs represent a versatile tool to activate transcription. We therefore sought to determine whether STARs and CRISPRi are functionally composable into network motifs that can dynamically control gene expression. To do this, we first demonstrated that STARs can be used to control expression of a single guide RNA (sgRNA) from a CRISPRi system^34^, in effect creating an activation-repression cascade (**Figure 6a** and **Supplementary Fig. 20**). As for the activation-activation cascade, an endoribonuclease RNA processing platform^48^ was used to cleave the transcriptionally fused target RNA-sgRNA transcript. By using a target RNA with low OFF level expression we were able to tightly control expression of the sgRNA in the absence of STAR, while achieving repression comparable to a constitutive sgRNA control when STAR was present (**Figure 6a**).

**Figure 6.**
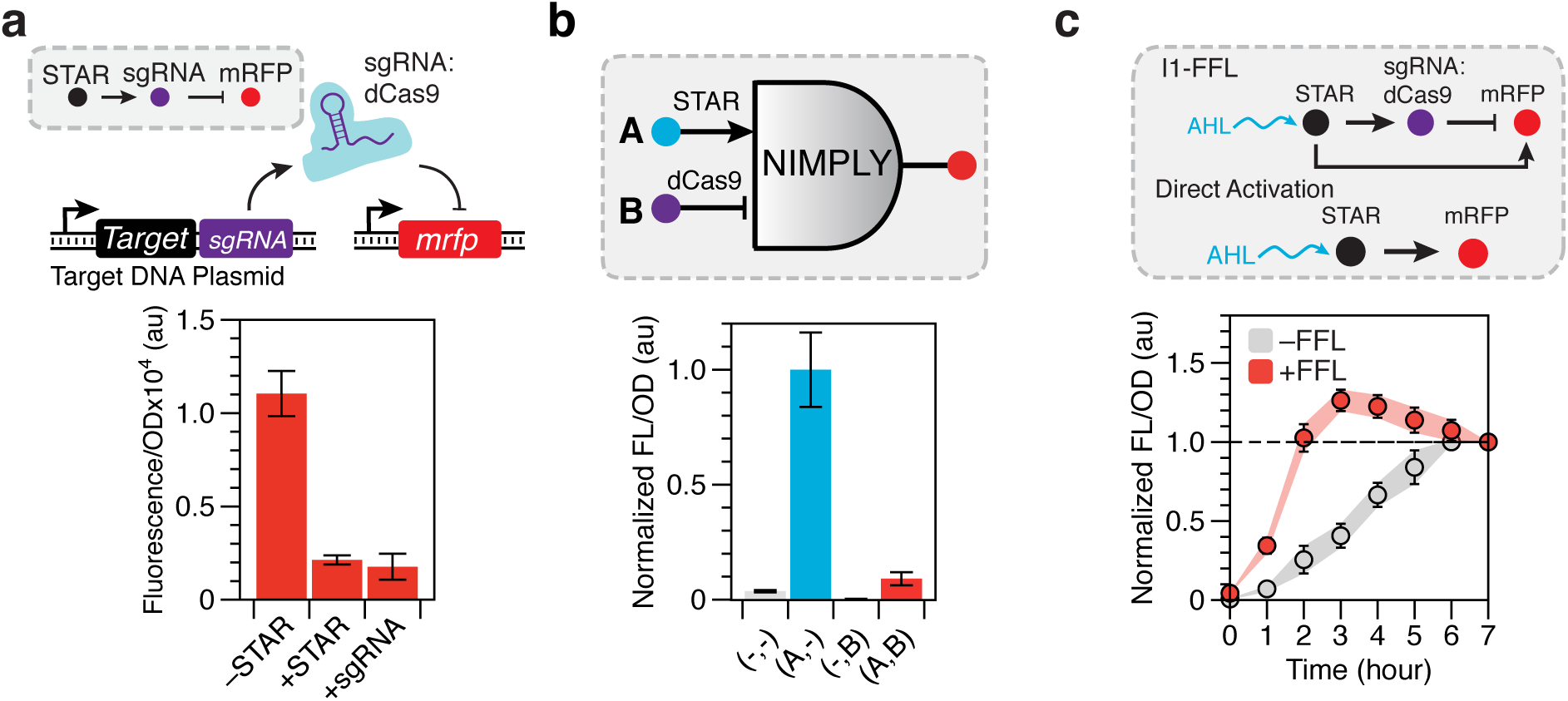
STARs and CRISPRi can be combined to create new types of dynamic genetic circuits. (**a**) STARs function with CRISPRi in an RNA activation-repression cascade. Schematic of the DNA template of a STAR regulated CRISPR interference (CRISPRi) cascade (see **Supplementary Fig. 20** for a detailed schematic). Fluorescence characterization was performed on *E. coli* cells transformed with a dCas9 and target RNA/mRFP expressing plasmid and a STAR-regulated single guide RNA (sgRNA) plasmid in the absence (-STAR) and presence (+STAR) of a plasmid encoding the cognate STAR. A constitutively expressed sgRNA (+sgRNA) was used as a repression control. Full controls used are shown in **Supplementary Fig. 20**. (**b**) STARs function with CRISPRi in logic gates. Schematic of a STAR and CRISPRi based NIMPLY (A AND NOT B) logic gate (see **Supplementary Figure 21** for a detailed schematic). Fluorescence characterization was performed on *E. coli* cells transformed with a plasmid encoding the NIMPLY logic gate and sgRNA in the presence of plasmids encoding all combinations of input A (STAR) and input B (dCas9). Fluorescence data were normalized to 1 for the ON condition (Input A only). (**c**) A STAR-CRISPRi incoherent type 1 feed-forward loop (I1-FFL) creates accelerated response and a pulse of gene expression. Schematic of the STAR-CRISPRi I1-FFL and direct activation control (see **Supplementary Fig. 21** for a detailed schematic). Fluorescence characterization of *E. coli* cells transformed with a plasmid encoding the I1-FFL (+FFL) or the direct activation (-FFL) cascade. STAR expression was induced at time 0 hour by addition of AHL and fluorescence measured every 1 hour for 7 hours. Repeats and no AHL controls are shown in **Supplementary Fig. 22**. Fluorescence data for each condition were individually normalized by dividing by the final fluorescence values at 7 hours for each colony before calculating the mean and s.d. at each time point. Fluorescence characterization was performed by bulk fluorescence measurements (measured in units of fluorescence [FL]/optical density [OD] at 600 nm). Data in (**a**, **b**) represents mean values of *n* = 9 biological replicas ± s.d and (**c**) is a representative of *n* = 3 biological replicas with repeats shown in **Supplementary Fig. 22**.

We next used the combination of activation and repression offered by both systems to create a NIMPLY logic gate (A AND NOT B) (**Supplementary Fig. 21**). Characterization of the NIMPLY gate revealed it to be functional – expressing mRFP only when the STAR (Input A) was present (**Figure 6b**).

Finally, the activation-repression cascade and the NIMPLY logic gate were combined to create the first RNA-based feed-forward loop^45^. The network architecture of an incoherent type 1 feed forward loop (I1-FFL) results in a temporal gene expression profile that shows accelerated response compared to direct activation, and an overshooting of the steady-state to create a pulse of gene expression^45^ (**Supplementary Fig. 21**). We observed both of these characteristics in our STAR and CRISPRi I1-FFL – with a response time (defined as time to reach half of the steady-state expression level) of 1.2 hours compared to 3.4 hours for direct activation, and a pulse of >25 % of the final steady-state expression level (**Figure 6c** and **Supplementary Fig. 22**).

Thus we have demonstrated that STARs and CRISPRi are highly composable, allowing for the creation of novel RNA-based network motifs. Given the numerous network motifs that require both activation and repression, the synergistic combination of high-performing CRISPRi and STARs provides a powerful toolset for the engineering of synthetic network motifs.

## Discussion

In this work, we have shown that by combining a simple RNA design motif with computational RNA structure design tools, we can easily design large libraries of high-performing RNA-based transcriptional activators called STARs. These new STARs achieve high functional performance, with the best-performing variants representing some of the highest dynamic ranges achieved by RNA-based regulators to date. Through analyzing the sequence-structure-function relationship of our STAR library, we uncovered design principles that we anticipate will further improve the robustness of our design strategy. In addition, we showed that orthogonal STARs can be identified entirely computationally, opening the door for large libraries of orthogonal and high-performing STARs to be engineered. Thus, the work described here represents a significant advancement of the STAR regulatory systems and our ability to computationally design RNA transcriptional regulators.

More broadly, STARs complement an increasingly comprehensive bacterial toolbox of powerful RNA-based regulators that are achieving protein-like dynamic ranges, such as the toehold translational activators^21^. Thus alongside CRISPRi transcriptional repressors^34^, STARs now offer a powerful RNA regulatory tool for controlling transcription. Interestingly, we observed that a distinct advantage of STARs over toehold switches was the tight control of gene expression in the OFF state, which in certain genetic contexts, gave rise to infinite folds of activation. We hypothesize that this reflects an important distinction between RNA transcriptional and translational activators. Both regulatory systems are dependent on a de-repressive mechanism – by default folding into RNA structures that prevent gene expression that are then de-repressed through binding of a *trans*-acting sRNA to achieve activation. As such, one of the key determinants of the level of gene expression in the OFF state is likely to be the misfolding of these RNA structures into non-repressive conformations. Toehold switch hairpins control translation, and are thus present for longer times during their regulatory regime. This longer time inherently allows for more RNA structural fluctuations that can be exploited by ribosomes to initiate translation even in the OFF state^50^. STARs on the other hand operate cotranscriptionally, meaning there is inherently a limited time-window in which these misfolding events can occur before the regulatory decision has been made. It is clear from our results that certain RNA sequences can fold into terminator structures extremely efficiently, which combined with the short time window of regulation leads to near perfect OFF levels. Of course this perfection in OFF level may incur a metabolic cost, as many rounds of abortive transcripts may be terminated before a STAR can capture one to activate it. Overall the comparison between toeholds switches and STARs reflects fascinating tradeoffs between equilibrium versus kinetic RNA design, and starts to uncover principles by which kinetically driven transient RNA structures may perform exquisite feats of regulation.

The emergence of high-performing RNA regulators has already seen their deployment in a variety of application spaces^32,33,51,52^. Contributing to this, we have demonstrated that STARs are ideally suited for a range of existing and novel applications. First, we found that STARs could be synergistically combined with existing regulators to create new composite systems for tunable control of gene expression. This included combining STARs with promoter-RBS strength variants to provide a new route to create switchable and tunable control elements from existing libraries^39^. In addition we showed that combining STARs with inducible promoter systems could tune their transfer functions, which to date has been largely achieved through engineering of DNA promoters and protein transcription factors^53^. Thus STARs add to an increasingly powerful toolbox of regulators that can be used in composite with existing regulatory mechanisms to alter their regulatory properties^54,55^.

We anticipate that the robustness of STAR regulation in diverse genetic and environmental contexts make them ideally suited for existing applications such as metabolic engineering. For example, the ability of STARs to regulate the expression of individual enzymes of metabolic pathways could be used to rapidly identify optimal expression regimes simply through titration of STAR expression level, circumventing the need to construct large libraries of pathway expression variants^56^. Moreover, the ability of STARs to regulate multigene metabolic pathways can provide new control points for pathway optimization, particular those that interfere with host metabolism and produce toxic intermediates or products. Finally, the relatively small size, functionality on the genome, and gene-independent regulation could make STARs ideally suited as a tool for strain engineering and reprogramming of cellular phenotypes. For example, STARs could be used to create synthetic switches for decoupling biomass and pathway production^57,58^.

STARs also add new tools for construction of transcriptional RNA-based network motifs. Importantly, these STARs not only add high-performing orthogonal transcriptional activators to the RNA genetic circuitry toolbox, but are functionally composable with CRISPRi transcriptional repressors, unlocking network motifs that require both activation and repression. As the repertoire of these basic network motifs increases, more complex composite motifs will be possible, allowing for more sophisticated gene control such as the creation of temporal regulatory programs^45^. As RNA synthetic biology begins to match the complexity of motifs achieved using protein-based transcription factors, we anticipate the emergence of several advantages of RNA-only transcriptional networks including smaller genetic footprint, potentially less cellular burden and faster network dynamics^2,24^.

Finally, one of the major advantages of RNA synthetic biology is the wealth of computational and experimental tools available to understand the sequence-structure-function relationship^2^. Interestingly, here we show that this can not only further design principles of synthetic RNA regulators but also potentially uncover principles of natural RNA regulators. For example, while there have been a wealth of studies examining how sequence and structure of terminator hairpins impacts terminator efficiency^26,59^, this work represents one of the few investigations into the effects of the 5’ sequence contexts of intrinsic terminators. Our results suggest that minimization of RNA structure upstream of a terminator hairpin can dramatically increase terminator efficiency. In agreement with this, single-molecule experiments have also shown that minimization of secondary structure upstream of terminator hairpins can increase terminator efficiency, suggesting terminator efficiency is not only an intrinsic property of the terminator hairpins but also of upstream sequence contexts^28^. As RNA synthetic biology begins to turn its attention to more diverse regulatory systems, we anticipate that through engineering we will uncover a deeper understanding of natural RNA regulators.

In summary, STARs represent a powerful class of RNA regulatory mechanisms that can be computationally designed to offer high-performing and orthogonal control of gene expression in diverse contexts. The uncovering of a kinetically driven RNA regulatory motif that can nonetheless be designed using equilibrium-focused computational design algorithms gives great hope for more advanced RNA synthetic biology. We anticipate that as we learn more about RNA cotranscriptional folding^27^, we will unlock even deeper principles of RNA design, and that the advances in RNA synthetic biology have only begun.

## Acknowledgements

The authors gratefully acknowledge the gift of TX-TL extract and buffers used in this work from Vincent Noireaux’s laboratory (University of Minnesota). The authors gratefully acknowledge the gift of dCas9 expression plasmids used in this work from Stanley Qi’s laboratory (University of Stanford). The authors gratefully acknowledge the gift of the deoxyviolacein expression plasmid from Robert Egbert of Adam Arkin’s laboratory (University of California, Berkley). The authors also thank John Dueber (University of California, Berkley) for helpful conversations about the deoxyviolacein pathway. The authors also thank Niles Pierce’s laboratory (California Institute of Technology) for helpful conversations about NUPACK and RNA design. This work was supported by an NSF CAREER Award [1452441 to J. B. L.].

## Methods

### Plasmid assembly

All plasmids used in this study can be found in **Supplementary Table 1** with key sequences provided in **Supplementary Tables 2-5**. STAR and target expressing DNA plasmids were constructed using inverse PCR (iPCR). Schematic of representative DNA plasmid maps used are shown in **Supplementary Fig. 3**. All assembled plasmids were verified using DNA sequencing.

### Integration of STAR regulated genes into the *E. coli* genome

Strains containing genomic insertions of STAR regulated genes were created using the clonetegration platform^60^ as summarized in **Supplementary Table 6**. The HK022 plasmid was used to integrate constructs into the *attB* site of the *E. coli* genome. Successful integrations were identified by antibiotic selection and colony PCR according to the published protocol^60^.

### Strains, growth media, *in vivo* bulk fluorescence measurements and *in vivo* flow cytometry fluorescence measurements

*In vivo* fluorescence characterization experiments were performed in *E. coli* strain TG1 except for **Supplementary Fig. 15** as described in the figure legend. For the in absence of STAR (-STAR) condition the no-STAR control plasmid (JBL002) was used. Experiments were performed for nine biological replicates collected over three separate days. It should be noted that for flow cytometry if cell counts did not meet the required threshold as described below because of slower growth rate, a fourth day of three more biological replicates was used. For each day of fluorescence measurements, plasmid combinations were transformed into chemically competent *E. coli* cells and plated on LB+Agar (Difco) plates containing combinations of 100 μg/mL carbenicillin, 34 μg/mL chloramphenicol and/or 50 μg/mL spectinomycin depending on plasmids used, and incubated approximately 17 hours (h) overnight at 37 °C. Plates were taken out of the incubator and left at room temperature for approximately 7 h. Three colonies were used to inoculate three cultures of 300 μL of LB containing antibiotics at the concentrations described above in a 2 mL 96-well block (Costar), and grown for approximately 17 h overnight at 37 °C at 1,000 rpm in a VorTemp 56 (Labnet) bench top shaker. Four μL of each overnight culture were then added to 196 μL (1:50 dilution) of either supplemented M9 minimal media (1 × M9 minimal salts, 1 mM thiamine hydrochloride, 0.4 % glycerol, 0.2 % casamino acids, 2 mM MgSO_4_, 0.1 mM CaCl_2_) or for the activation-activation cascade and CRISPRi experiments (**Figure 5b**, **6a-b** and **Supplementary Fig. 20**) MOPS EZ rich defined media (Teknova) (1 × MOPS mixture, 1.32 mM K_2_HPO_4_, 1 × ACGU, 1 × supplement EZ, 0.2 % glucose) containing the selective antibiotics and grown for 6 h at the same conditions as the overnight culture. For **Figure 6a,b** and **Supplementary Fig. 20** 4 μL of 5 μg/mL of anhydrotetracycline (aTc) was also added (1:50 dilution). For **Figure 2b** and **Supplementary Fig. 14** 4 μL of acyl-homoserine lactone (AHL) (N-(β-ketocaproyl)-L-Homoserine lactone, Cayman Chemical) solutions was also added (1:50 dilution). For **Figure 6c** and **Supplementary Fig. 22** 16 μL of each overnight culture were added to 768 μL of MOPS EZ rich defined media containing selective antibiotics and grown for 4 h at the same conditions as the overnight culture. After 4 h 16 μL of either 5 μM of AHL or ddH_2_0 were added and cultures grown for 7 h and fluorescence measured every 1 hour. For all bulk fluorescence measurements after subculture 50 μL of this culture was then transferred to a 96-well plate (Costar) containing 50 μL of phosphate buffered saline (PBS). Fluorescence (FL) and optical density (OD) at 600 nm were then measured using a SynergyH1 plate reader (Biotek). The following settings were used: sfGFP fluorescence (485 nm excitation, 520 nm emission), mRFP fluorescence (560 nm excitation, 630 nm emission) or YFP fluorescence (510 nm excitation, 540 nm emission).

For *in vivo* flow cytometry measurements for **Figure 1b, d**, **Figure 5d**, **Supplementary Fig. 4 Supplementary Fig.11** and **Supplementary Fig. 15**, subcultures (6-12 μL) were transferred into a FACS round-bottom 96 well plate with 244 μL of PBS containing 2mg/mL kanamycin to stop translation. The plate was then read on a BD LSR II using the high-throughput setting with the high-throughput sampler (HTS). The following parameters were collected on the BD LSR II: forward scatter (FSC), side scatter (SSC), and sfGFP fluorescence (488 nm excitation, 515-545 nm emission). Each sample was required to have at least 5,000 counts or up to 50,000 counts. Counts were gated as explained in **Supplementary Note 4**. sfGFP fluorescence values were recorded in relative channel number (1-262,144 corresponding to 18-bit data) and the geometric mean over the gated data was calculated for each sample.

For *in vivo* flow cytometry measurements for **Figure 1c** and **Supplementary Fig. 6**, subcultures (1-12 μL) were transferred into a FACS round-bottom 96 well plate with 250 μL of PBS containing 2mg/mL kanamycin to stop translation. The plate was then read on a BD Accuri C6 Plus CSampler. The following parameters were collected on the BD Accuri C6 Plus: forward scatter (FSC), side scatter (SSC), and sfGFP fluorescence (FL1: FITC 488 nm excitation, 518-548 nm emission). Each sample was measured up to 50,000 counts. Counts were gated as explained in **Supplementary Note 4**. sfGFP fluorescence values were recorded in relative channel number (1-16,777,216 corresponding to 24-bit data) and the geometric mean over the gated data was calculated for each sample.

### Bulk fluorescence data analysis

On each 96-well block there were two sets of controls; a media blank and *E. coli* TG1 cells transformed with combinations of control plasmids JBL001, JBL002 or JBL5999 (blank cells) and thus not expressing reporter genes (**Supplementary Table 1** and **Supplementary Fig. 3**). The block contained three replicates of each control. OD and FL values for each colony were first corrected by subtracting the corresponding mean values of the media blank. The ratio of FL to OD (FL/OD) was then calculated for each well (grown from a single colony) and the mean FL/OD of blank cells was subtracted from each colony’s FL/OD value. Three biological replicates were collected from independent transformations, with three colonies characterized per transformation (9 colonies total). Means of FL/OD were calculated over replicates and error bars represent standard deviations (s.d). Normalized FL/OD was calculated by dividing the mean values of a specific condition indicated in the figure legend and propagating errors. For **Figure 6c** and **Supplementary Fig. 22** fluorescence data for each condition were individually normalized by dividing by the final fluorescence values at 7 h for each colony before calculating the mean and standard deviation at each time point.

### Flow cytometry data analysis

Spherotech 8-Peak Rainbow Calibration Beads (Spherotech) were used to obtain a calibration curve to convert fluorescence intensity (geometric mean, relative channel number) into Molecules of Equivalent of Fluorescein (MEFL). For each experiment, *E. coli* cells transformed with combinations of control plasmids JBL001 and JBL002 (blank cells) not expressing sfGFP were used as a control. The mean MEFL value of blank cells without sfGFP expression was subtracted from each colony’s MEFL value. Mean MEFL values were calculated for at least 7 biological replicates and error bars represent the standard deviation (s.d). The ON level (+STAR) is the MEFL of cells containing the target expressing DNA plasmid and the STAR expressing plasmid and the OFF level (-STAR) is the MEFL of cells containing the target expressing DNA plasmid and a no-STAR control plasmid (JBL002). The fold activation was calculated by dividing the ON level by the OFF level (ON/OFF). For **Figure 1b** a Welch’s t-test was performed on each -STAR/+STAR condition with all STAR variants showing statistically significant differences between -STAR and +STAR conditions (*P* < 0.05).

### *In vivo* catechol 2,3-dioxygenase and deoxyviolacein pathway characterization and data analysis

For *in vivo* catechol 2,3-dioxygenase absorbance measurements *E. coli* strain TG1 was used. Plasmids were transformed, grown overnight and subculture as described for *in vivo* bulk fluorescence measurements. After subculture growth, OD at 600 nm was measured and cells diluted to an OD at 600 nm of 0.1 in 100 μL of M9 media. 98 μL of these cells were then transferred to a 96-well plate (Costar) and 2 μL of 25 mM catechol (Sigma Aldrich) added. Conversion of catechol to 2-hydroxymuconate semialdehyde was characterized by measuring OD at 385 nm in a BioTek SynergyH1 microplate reader for 25 minutes at 5 minute intervals and reactions held at 33 °C. The maximum rate of catechol to 2-hydroxymuconate semialdehyde conversion was determined between 0 and 10 minutes as described in **Supplementary Fig. 18**. Repeats and controls were as described for bulk fluorescence measurements. For *in vivo* deoxyviolacein quantification *E. coli* strain DH5 alpha pir used. Plasmids were transformed, grown overnight as described for *in vivo* bulk fluorescence measurements. 150 μL of overnight culture was then removed and pelleted by centrifugation at 15,000 rpm for 5 minutes. The pellet was resuspended in 150 μL of 100 % ethanol and cells lysed by boiling for 5 minutes, followed by centrifugation at 15,000 rpm for 5 minutes. One hundred μL of supernatant was then carefully removed, added to a costar 96-well plate and the deoxyviolacein quantified by measuring OD at 525 nm in a BioTek SynergyH1 microplate reader.

### Total RNA extraction and DNase treatment for reverse transcription quantitative PCR (RT-qPCR)

For extraction of total RNA for RT-qPCR experiments *E. coli* strain TG1 was used. Plasmids were transformed and subsequent colonies grown overnight as described for *in vivo* bulk fluorescence measurements. For each biological replicate, 20 μL of a single overnight culture was added to three wells containing 980 μL (1:50 dilution) of supplemented M9 minimal media containing the selective antibiotics and grown for 6 h at the same conditions as the overnight cultures. For each plasmid combination, 500 µL of cells were removed from three wells (grown from one colony) and combined into a 1.6 mL tube and pelleted by centrifugation at 13,000 rpm for 1 min. Total RNA was extracted and DNase treated according to a previously described protocol^30^.

### Normalization of total RNA, reverse transcription and qPCR measurements

To enable comparison between different samples, each DNase treated sample was normalized to contain the same total RNA concentration. Each sample was quantified using a Qubit Fluorometer and the sample diluted to 0.025 ng/µL of total RNA in 20 µL RNase free ddH_2_0. For cDNA synthesis 1 µL of this total RNA, 1 µL of 2 µM reverse transcription primer (**Supplementary Table 7**), 1 µL of 10 mM dNTPs (New England Biolabs) and RNase free ddH_2_0 up to 6.5 µL were incubated for 5 min at 65 °C and cooled on ice for 5 min. 0.25 µL of Superscript III reverse transcriptase (Life technologies), 1 µL of 100 mM Dithiothreitol (DTT) and 1x first-strand buffer (Life technologies), 0.5 µL RNaseOUT (Life Technologies) and RNase free H_2_O up to 3.5 µL were then added, incubated at 55 °C for 1 h, 75 °C for 15 min and then stored at -20 °C. qPCR was performed using 5 µL of Maxima SYBR green qPCR master mix (Thermo Scientific), 1 µL of cDNA solution and 0.5 µL of 2 µM sfGFP qPCR primers (**Supplementary Table 7**) and RNase free ddH_2_O up to 10 µL. A ViiA 7 real-time PCR machine (Applied Biosystems) was used for data collection using the following PCR program: 50 °C for 2 min, 95 °C for 10 min followed by 30 cycles of, 95 °C for 15 sec and 60 °C for 1 min. All measurements were followed by melting curve analysis. A MicroAmp EnduraPlate Optical 384-well plate (Applied Biosystems) and an optically clear seal (Applied Biosystems) were used for all measurements. Results were analyzed using the ViiA 7 software (Applied Biosystems) by a relative standard curve. For quantification, a five point standard curve covering a 10000-fold range of sfGFP DNA concentrations was run in parallel and used to determine the relative sfGFP cDNA abundance in each sample. It was shown that the sfGFP qPCR primer set had a primer efficiency ∼80%. All cDNA samples were measured in triplicate and non-template controls run in parallel to control for contamination and non-specific amplification or primer dimers. In addition, qPCR was performed on total RNA samples to confirm no DNA plasmid was detected under the conditions used. Melting curve analysis and DNA gel electrophoresis were performed to confirm that only a single product was amplified.

### Characterization of *E. coli* motility

Plasmids were transformed and subsequent colonies grown overnight as described for *in vivo* bulk fluorescence measurements in *E. coli* strains described in figure legends. For each biological replicate, 40 μL of a single overnight culture was added to a culture tube containing 1960 μL (1:50 dilution) of LB containing the selective antibiotics and grown for 6 h at the same conditions as the overnight cultures. We note that 17 μg/mL chloramphenicol for the genomically integrated chloramphenicol resistance cassettes was used. After subculture, OD at 600 nm was measured and cultures diluted to an OD at 600 nm of 0.250 with fresh LB containing selective antibiotics. 3 µL of diluted cultures were plated on to the center of 0.25% tryptone agar (1 % tryptone, 0.5 % NaCl, 0.25 % agar) plates (each containing 25 ml of 0.25% tryptone agar) containing selective antibiotics, air dried for 10 minutes and incubated overnight at 30 °C on a flat surface. Plates were imaged under trans white light illumination using a Chemidoc XRS+ (BioRad).

### Characterization of STARs in TX-TL

TX-TL cell extract, amino acids and energy solution were gifted by the Noireaux lab and prepared as described in Garamella *et al*^40^. Extract, amino acids and energy solution and a buffer (containing Mg-glutamate, K-glutamate and PEG-8000) were combined and pre-incubated on a heat block at 37 °C for 20 min. After incubation 7.88 µL of the TX-TL mix was added to 2.63 µL of DNA and water solution. Each reaction contained 8 nM of a target DNA plasmid and either 15 nM of a no-STAR control plasmid or 15 nM of a DNA plasmid encoding a cognate STAR. Dilutions of DNA plasmids were calculated using a modified spreadsheet derived from Sun *et al*^41^. For TX-TL characterization of STAR-regulated catechol 2,3-dioxygenase, 1 mM of catechol (Sigma Aldrich) was added while maintaining a total volume of 10.5 µL. 10 µL of each reaction was then placed into their respective wells in a 384-well plate (Nunc) and sealed with an optical clear seal (Thermo Scientific) and measured in a BioTek SynergyH1 microplate reader. For fluorescence measurements, reactions were held at 29 °C and measured every 5 minutes (485 nm excitation, 520 nm emission). For catechol 2,3-dioxygenase expression reactions were held at 33 °C and OD at 385 nm measured every 5 minutes. On each 384-well plate a control consisting of a TX-TL reaction containing no DNA was used and the FL or OD values of this reaction were subtracted from each condition to remove background signal from the TX-TL reaction. Rate of sfGFP production was determined by taking the time derivative of fluorescence measurements, and the average rate of sfGFP production calculated from the linear sfGFP production phase as described in **Supplementary Fig. 17**. *n* = 3 biological replicas were collected for each independent repeat, for a total of three independent repeats. Mean values were calculated from these *n* = 9 replicas and error bars represent standard deviations (s.d).

